# Computationally efficient assembly of a *Pseudomonas aeruginos*a gene expression compendium

**DOI:** 10.1101/2022.01.24.477642

**Authors:** Georgia Doing, Alexandra J. Lee, Samuel L. Neff, Jacob D. Holt, Bruce A. Stanton, Casey S. Greene, Deborah A. Hogan

## Abstract

Over the past two decades, thousands of RNA sequencing (RNA-seq) gene expression profiles of *Pseudomonas aeruginosa* have been made publicly available via the National Center for Biotechnology Information (NCBI) Sequence Read Archive (SRA). In the work we present here, we draw on over 2,300 *P. aeruginosa* transcriptomes from hundreds of studies performed by over seventy-five different research groups. We first developed a pipeline, using the Salmon pseudo-aligner and two different *P. aeruginosa* reference genomes (strains PAO1 and PA14), that transformed raw sequence data into a uniformly processed data in the form of sample-wise normalized counts. In this workflow, *P. aeruginosa* RNA-seq data are filtered using technically and biologically driven criteria with characteristics tailored to bacterial gene expression and that account for the effects of alignment to different reference genomes. The filtered data are then normalized to enable cross experiment comparisons. Finally, annotations are programmatically collected for those samples with sufficient meta-data and expression-based metrics are used to further enhance strain assignment for each sample. Our processing and quality control methods provide a scalable framework for taking full advantage of the troves of biological information hibernating in the depths of microbial gene expression data. The re-analysis of these data in aggregate is a powerful approach for hypothesis generation and testing, and this approach can be applied to transcriptome datasets in other species.

**Significance:** *Pseudomonas aeruginosa* causes a wide range of infections including chronic infections associated with cystic fibrosis. *P. aeruginosa* infections are difficult to treat and people with CF-associated *P. aeruginosa* infections often have poor clinical outcomes. To aid the study of this important pathogen, we developed a methodology that facilitates analyses across experiments, strains, and conditions. We aligned, filtered for quality and normalized thousands of *P. aeruginosa* RNA-seq gene expression profiles that were publicly available via the National Center for Biotechnology Information (NCBI) Sequence Read Archive (SRA). The workflow that we present can be efficiently scaled to incorporate new data and applied to the analysis of other species.

## Introduction

The opportunistic pathogen *Pseudomonas aeruginosa* is found in a wide array of environments. *P. aeruginosa* is found in soil (1) and freshwater samples (2, 3), and it is cultured for biotechnology applications (4, 5). It causes diverse infections including those common in chronic lung infections of people with cystic fibrosis (CF) where it is difficult to eradicate (6). The factors that lead to its persistence are not fully understood. Analyses of *P. aeruginosa* isolates, including many clinical isolates, show significant diversity both in genome content and in recently evolved genetic changes. In addition to genomic variation, the versatility of *P. aeruginosa* is also attributable to behavioral adaptive changes driven by gene expression. The breadth of *P. aeruginosa* research is reflected in the abundance of publicly available transcriptional datasets from both microarray (∼1,000 profiles) and RNA-seq technologies (∼3,000 profiles). Across multiple areas of interest, many studies of *P. aeruginosa* have examined clinical isolates and the laboratory strains such as PAO1 and PA14, which have served as models of transcriptional regulation. Given the unusually high number of transcription factors, sigma factors and two component systems (7–9), transcriptional profiling across conditions and mutant genotypes has been a fruitful approach to better understand *P. aeruginosa* physiology.

The *P. aeruginosa* community has long supported the development and widespread use of databases, hubs and analysis tools, such as The *Pseudomonas* Genome Database (10), BACTOME (11), the International *Pseudomonas* Consortium Database (12), the *Pseudomonas aeruginosa* metabolome database (13), the *Pseudomonas aeruginosa* transcriptome viewer (14), and the shiny app with algorithmically annotated datasets, GAPE (15). Tools have also been developed that utilize public data from across many experiments in concert, such as the ADAGE web server which enables the exploration of *P. aeruginosa* microarray data after processing by a machine learning algorithm (16). To further support the development of resources that leverage public RNA-seq data, we present a computationally efficient method to reprocess reads from RNA-seq datasets (**Figure 1** for overview). After validating our methods for high throughput mapping of reads, we collected publicly available *P. aeruginosa* RNA-seq data, processed it, filtered samples that did not meet quality control metrics, then normalized the data (**Figure 1**, steps 1-4). We assessed this approach by examining correlations between co-regulated genes and using strain-specific markers (**Figure 1**, steps 5-6). Lastly, we summarized the meta-data annotations to provide users with information on the composition of the compendium (**Figure 1**, steps 7-8). Thus, we present a method to build a uniform compendium made from public data and demonstrate its success using *P. aeruginosa* gene expression profiles.

**Figure 1.**
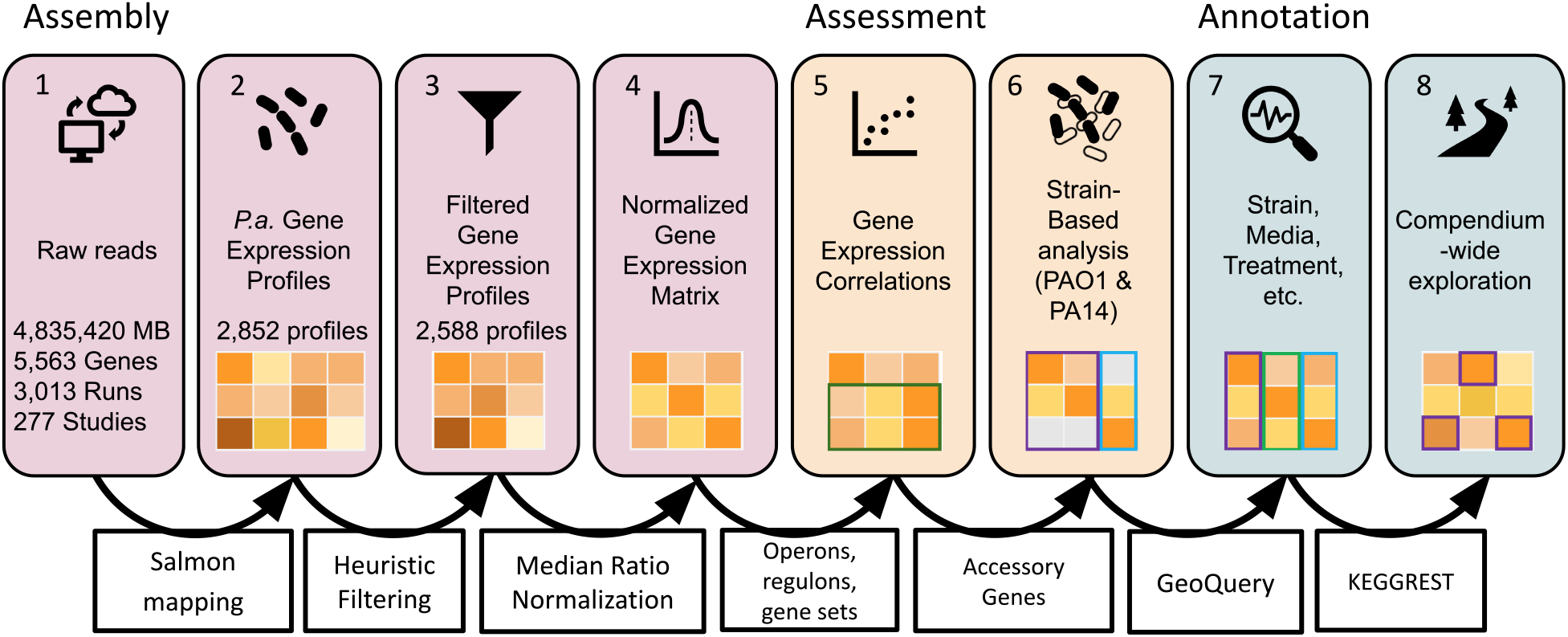
The steps and corresponding methods (open boxes attached to arrows) for assembly (1-4), assessment (5-6) and annotation (7-8) of a uniform compendium of public gene expression profiles from RNA-seq data of *P. aeruginosa* (*P*.*a*.). **1)** Raw reads from the Sequence Read Archive (SRA) totaled over 4 million megabases (MB) from an organism with approximately 5,563 genes. “Runs” refer to one or more fastq files containing the reads from the sequencing of a single RNA sample and “Studies” refer to sets of Runs deposited together. **2)** *P*.*a*. gene expression profiles refer to the results of aligning reads from a sample (referred to in SRA as “Experiment”) to a reference genome and can be read counts or transcripts per million. **3)** Profiles were filtered to remove those that did not meet empirically determined quality criteria. **4)** Profiles were normalized using the Median of Ratios method. **5)** Sets of co-regulated genes were used to benchmark target patterns. **6)** The compendium was assessed by strain-specific gene expression patterns. **7)** The compendium was annotated for strain, media, genetic modifications, treatment and other fields of interest. **8)** Pathway and function data facilitated compendium-wide exploration.

## Results & Discussion

### Assembly of a *P. aeruginosa* gene expression compendium

A total of 3,013 NCBI “Run” accession numbers, organized into 2,867 samples (indexed by SRA “Experiment” accession numbers), were used to query and download all associated fastq files. For each of the 2,867 samples, all associated Runs (fastq files) were downloaded and mapped, using Salmon, to transcriptional indices built from PAO1 and PA14 cDNA libraries. Samples had multiple associated Run files for multiple reasons including paired end read libraries. Gene expression profiles from fastq files mapped to each reference were assembled into separate compendia of 2,852 profiles derived from the same set of 2,852 samples that were successfully downloaded and mapped to both references.

### Salmon mapping produces similar transcriptional profiles of *P. aeruginosa* to field-standard alignment

Pseudo-alignment algorithms estimate transcriptional profiles from high-throughput sequencing reads in a fraction of the time required by traditional alignment algorithms and make the re-processing of thousands of RNA-seq datasets practical (17, 18). Pseudo-alignment has been thoroughly validated and widely used on eukaryotic RNA-seq data (19, 20). However, these algorithms are used less widely in microbial research. As a first step in the creation of a normalized compendium of publicly available *P. aeruginosa* RNA-seq data, we validate Salmon pseudo-alignment (mapping) for the analysis of *P. aeruginosa* gene expression data.

To assess the results of RNA-seq data mapping using the pseudo-aligner Salmon (18), we compared its output to the results of field-standard alignment using the CLC Genomics Workbench (CLC). For this, we performed RNA-seq on samples that have well-characterized transcriptional differences: *P. aeruginosa* strain PA14 wild type and a *pstB*::Tn*M* mutant derivative grown as colony biofilms on minimal medium. We chose this comparison because *pstB* mutants have a constitutively active low-phosphate response (21), which is a transcriptionally driven process (22) that would give a clear differential expression signal. Each of the wild-type and *pstB* mutant samples was processed using both Salmon and CLC.

An important parameter for Salmon pseudo-alignment is the “library type,” which is determined by whether the reads were obtained via paired end or single end (unpaired) sequencing. Publicly available *P. aeruginosa* RNA-seq data consists of a mix of paired and unpaired reads. The libraries from the wild-type and *pstB*::Tn*M* samples contained paired end reads, and we compared the results of mapping with the library type flag set to “paired” and “unpaired”. When the data were mapped specifying the library type as “paired”, Salmon had lower estimates than CLC for the expression of many genes, especially those that were lowly expressed, and this difference was not observed when the library type was specified as “unpaired” (**Supplemental Figure S1**). Linear models for the comparison of CLC- and Salmon-generated transcripts per million (TPM) as “paired” (average R^2^ across samples of 0.68) showed a lower fit than that between CLC and Salmon-generated data generated as “unpaired” (average R^2^ of 0.78).

We suspected that differences in TPM values between Salmon and CLC were due to the polycistronic nature of the *P. aeruginosa* genome, which violates assumptions made by Salmon for paired end libraries. Pseudo-alignment algorithms pre-compute transcriptome indices for k-mer mapping of reads, mapping only to coding regions of a genome. When genes are in operons, the assumption that forward and reverse reads (each 50-150 bp long) would map to the same transcriptome index (same gene), does not necessarily hold. To determine if the improved concordance between CLC and Salmon with unpaired mode was due to correction for polycistrons, we analyzed the correlations separately for monocistronic genes and polycistronic operons. We found that both mono- and polycistronic operons had better fits between CLC- and Salmon-generated data when processed as unpaired (R^2^ of 0.89 and 0.91, respectively) compared to when processed as paired (R^2^ of 0.86 and 0.83, respectively). These data suggested that while treating the data as paired resulted in slight differences in concordance dependent on operon size, treating the data as unpaired improved concordance for all genes regardless of operon size. Therefore, we maintained the parameter of “unpaired” reads for all datasets regardless of the library type.

### Salmon mapping and CLC alignment produce similar differential expression analysis

To further evaluate Salmon for the analysis of *P. aeruginosa* RNA-seq data, we performed differential expression (DE) analyses, comparing the expression profiles of *P. aeruginosa* strain PA14 wild type to a *pstB*::Tn*M* mutant for the expected difference in the PhoB-controlled low phosphate response (21). Alignment by CLC and mapping by Salmon to the strain PA14 reference generated read count values per gene that were highly similar for the two mapping methods for the datasets of WT and *pstB*::Tn*M P. aeruginosa* (**Figure 2A**, left axis and **Supplemental Dataset S1**). One gene, *pqqA*, which is a short gene consisting of only 72 nucleotides, appeared to have higher counts as estimated by Salmon than CLC (**Figure 2A**). The overall variation between the alignment methods (R^2^ = 0.99) (**Figure 2A**, left axis) was much less than that identified between the two genotypes of *P. aeruginosa*, wild type and *pstB*::Tn*M* (R^2^ = 0.72) (**Figure 2A**, right axis). Genes regulated by PhoB (indicated in **Figure 2A** by large circle size) had higher expression in *pstB*::Tn*M* than the wild type, with consistent expression across alignment methods that suggested an upregulation of the low-phosphate response in the *pstB* mutant, as expected.

**Figure 2.**
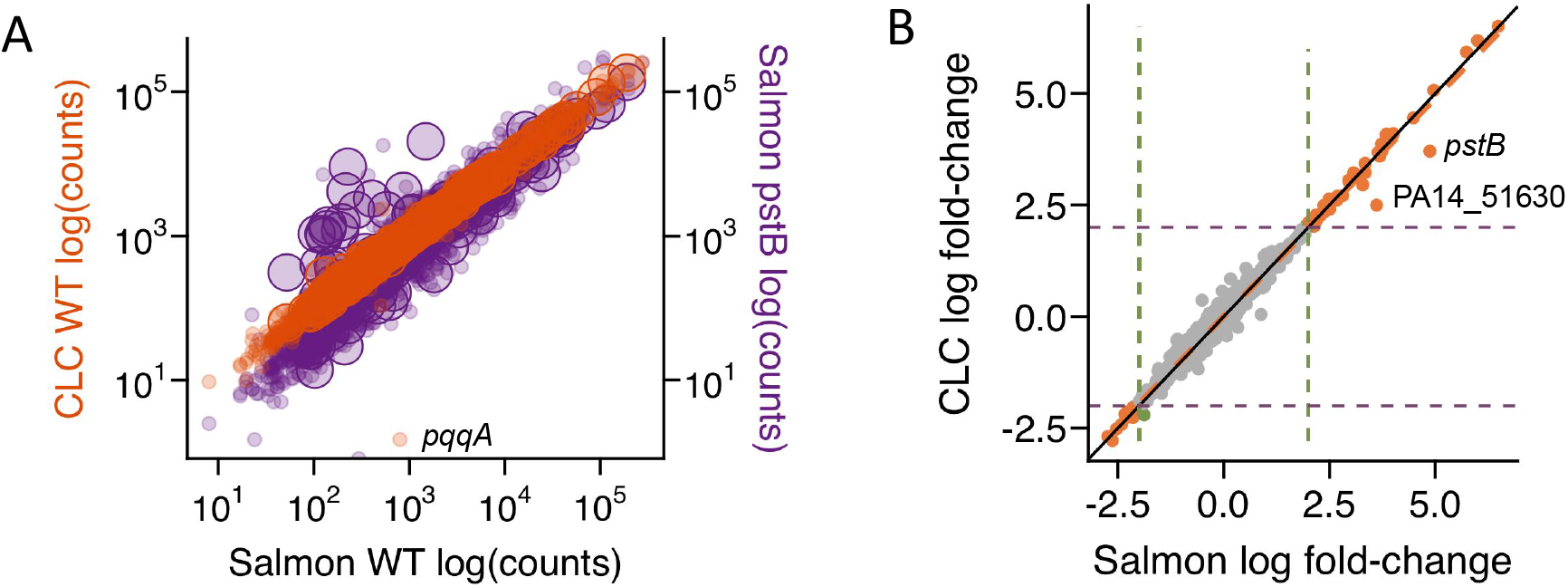
Validation of the Salmon pseudo-alignment method for the analysis of *P. aeruginosa* RNA-seq data. **A)** Correlations of log_10_ raw counts for *P. aeruginosa* wild type grown in MOPS minimal medium with 0.7 mM phosphate (n = 2) as determined by Salmon were highly similar to those determined by CLC (orange). The variation in the data across the two methods was less than the difference between the wild type and *pstB*::Tn*M*, grown in the same conditions (n = 2), when all samples were analyzed by Salmon (purple). PhoB-regulated genes, which were expected to be higher in *pstB*::Tn*M*, are indicated with larger circles. **B)** Fold-change values between wild-type and *pstB*::Tn*M* strain data were determined for both Salmon or CLC. Points with an absolute value log_2_ fold-change >2 by both methods are indicated by orange points. Three genes (PA14_07380, PA14_19680 and PA14_62690) had absolute value log_2_ fold-change > 2 by CLC but not Salmon (three green points that appear overlapping).

Differential expression analysis between WT and *pstB*:Tn*M* using EdgeR (23) produced similar per-gene fold-change values with data from CLC and Salmon (R^2^ = 0.98) (**Figure 2B** and **Supplemental Dataset S1**). Differentially expressed genes identified by both methods included genes that we expected to be induced in the *pstB* mutant including the genes encoding a histidine kinase sensor (*phoR)*, a well-studied alkaline phosphatase (*phoA)*, a periplasmic phosphate sensing appendage (*pstS)*, machinery for the import of both phosphate and phosphonate (*phnC*), secreted phosphate scavengers such as phospholipase C (*plcN*) and an extracellular DNase (*eddA)* as well as an extracytoplasmic sigma factor (ECF) known to interact with PhoB in the RNA polymerase holoenzyme (*vreI)* (**Supplemental Dataset S1**) (22). All of these genes were significantly different in both Salmon- and CLC-based analyses. There were differences between the statistical scores produced by the differential expression analyses, (R^2^ = 0.81), though much of the variation came from differences in genes with relatively high significance scores for differential expression in both alignments (adjusted p > 0.05). Since significance scores are often interpreted based on thresholds (i.e., adjusted p < 0.05), variation in significance scores well above or below this threshold is unlikely to influence biological interpretations.

CLC identified 70 genes with significance scores below the common threshold of adjusted p > 0.05 that Salmon did not (**Supplemental Dataset S1**), which suggests that significance values may require adjustment for hypothesis generation depending on the method of read alignment used. However, none of these genes had log_2_ fold-change magnitudes greater than two and thus none would be considered differentially expressed genes, regardless of their statistical score if the commonly used two-fold change metric (absolute value log_2_ fold-change > 2) was also used as a criterion. Two genes, while they met the standard fold-change cutoff by both methods, were mapped at lower levels by CLC compared to Salmon, including *pstB* (PA14_70810, PA5366), whose mapping or alignment may have been affected by the transposon insertion, and PA14_51630 (PA0978), encoding a hypothetical protein (**Figure 2B**). Three other genes had statistically significant and sufficient magnitude log_2_ fold-change values by CLC-aligned data, but not Salmon-mapped data, and they encoded hypothetical proteins PA14_07380 (PA0568), PA14_19680 (PA3432) and PA14_62690 (PA4739). These genes had marginally different fold-changes that fell very close to the cutoff for DEGs. The reasons for why a small number of genes showed differential results upon analysis by different methods are not known, but these data underscore the need for independent validation approaches when identifying candidates for further study.

### Salmon is robust to reference genome strain

The genomes of thousands of environmental and clinical isolates of *P. aeruginosa* have been sequenced and revealed a population structure that consists of multiple clades including the distinct clades that contain strain PAO1 and PA14 (10). Isolates can differ from each other by thousands of SNPs and hundreds of accessory genes. Numerous strains of *P. aeruginosa* have been analyzed by RNA-seq, and there is value in being able to leverage the large collection of genetically diverse data by mapping reads from multiple strains to a single reference genome for speed and characterization using a common set of genes.

To determine the impact of analyzing RNA-seq data using a reference genome from a different strain by Salmon, we compared the analysis of our strain PA14 RNA-seq data using both the PA14 reference genome (assembly ASM1462v1) and the reference genome for strain PAO1 (assembly ASM676v1), a distinct strain from a different phylogenetic clade. Most of the data in the compendium was generated using one of these two strains. We found that Salmon results for PAO1 and PA14 homologs were highly similar regardless of whether a PAO1 or PA14 reference genome was used (average R^2^ = 0.93, range of 0.91-0.94 for log_10_ counts across all four samples) (**Figure 3A**), which is consistent with the fact that over 90% of genes in PAO1 and PA14 strains have high homology (23, 24). In differential expression analyses between wild-type and *pstB*::Tn*M* strains, using Salmon-aligned data with PAO1 and PA14 reference genomes, only two genes, those from a single operon PA14_51620 (PA0978) and PA14_51630 (PA0979), both encoding hypothetical proteins had high magnitude fold-change values when aligned to PA14 but not when aligned to PAO1 (**Figure 3B**). It is worth noting that one of these loci also showed discrepancies between CLC and Salmon alignment. These PAO1 and PA14 homolog pairs had 95.5% and 96.7% identity, which is lower than average for PAO1 and PA14 homologs (99.1% identity).

**Figure 3.**
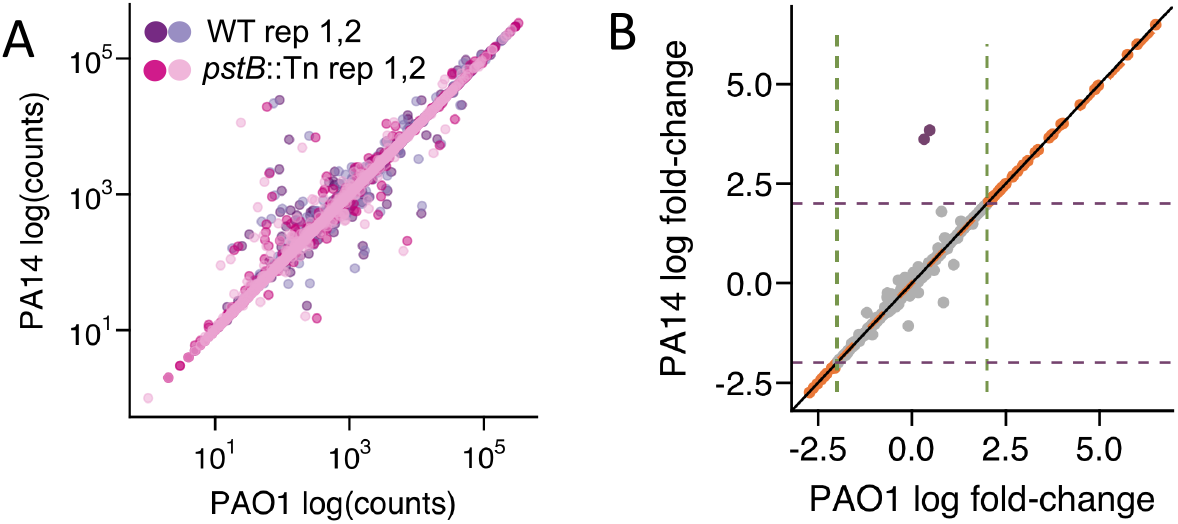
Analysis of the effects of reference genome on expression analysis. **A)** Mapping of *P. aeruginosa* strains PA14 WT and *pstB*::Tn*M* reads (log_10_ counts) using PAO1- or PA14-based transcriptional indices showed that some genes had more reads for the PA14 genome while other genes had more reads for the PAO1 genome; the same genes showed methodology-dependent differences in mapping across samples. **B)** Most high magnitude values, with log_2_ fold-change > 2, derived from PAO1-aligned data were highly similar to those from PA14-aligned data (orange) and two genes had high magnitude fold-change values by PA14-aligned, but not by PAO1-aligned data (purple).

### Heuristic-based filtering selects gene expression profiles with comparable distributions

Analysis of the *P. aeruginosa* RNA-seq compendia suggested that there were a number of samples that did not meet our quality score cut-offs for inclusion in these particular compendia. To analyze this problem, we focused on two profile characteristics within the compendia of data that provided heuristics for quality control including sparsity (the number of genes with zero counts representing undetected transcripts) and the median expression of a set of ten housekeeping genes (HKs) (*ppiD, rpoD, rpoS, proC, recA, rpsL, rho, oprL, tipA* and *nadB*). Heuristics were determined separately for compendia constructed by alignment of data to either strain PAO1 or PA14. We chose this approach in order to retain samples that might have reduced mapping rates due to the use of a reference genome different from that of the strain used. Strain-specific filter values for both of these parameters were empirically determined from 911 profiles of PAO1 samples aligned to a PAO1 reference and 588 profiles of PA14 samples aligned to a PA14 reference; these subsets of profiles were used because the strain identity was annotated in the sample record. Using these subsets with known strain identities, the thresholds for sparsity were chosen to be at the 10^th^ and 90^th^ percentiles. The PAO1 range for the zero-count genes per sample was 8 - 1,037 and the PA14 range was 10 - 1,037. In **Figure 4**, the sparsity values for profiles kept and removed are shown. Removed profiles had too many genes without any detected transcripts (**Figure 4A**, orange symbols). The thresholds for median housekeeping gene expression were the 20^th^ and 98^th^ percentiles (PAO1 HKs: 211.0 – 839.5 TPM, PA14 HKs: 258.4 - 840.7 TPM) (**Figure 4B**, orange symbols). Samples were included in the compendium if they passed both sparsity and housekeeping gene heuristics upon analysis using either the PAO1- or PA14-mapped data. Of the 2,852 samples processed, 2,333 samples were retained after filtering (**Supplemental Dataset S2**). Of the 519 samples removed, 41% were from PAO1 samples, 19% were from PA14 samples, 15% were clinical isolates, and 24% lacked strain annotation. There was not one particular strain type that failed in these criteria. Many of the samples that did not meet these criteria were from experiments generated by ribosomal profiling, RNA immunoprecipitation sequencing (RIP-seq) or Global sRNA Target Identification by Ligation and Sequencing. Any other samples that were generated using these methodologies were also removed from the compendium. Because the heuristic-based filtration criteria are not direct measures of technical quality of a sample, filtering cut-offs should be adjusted and optimized for different downstream uses.

**Figure 4.**
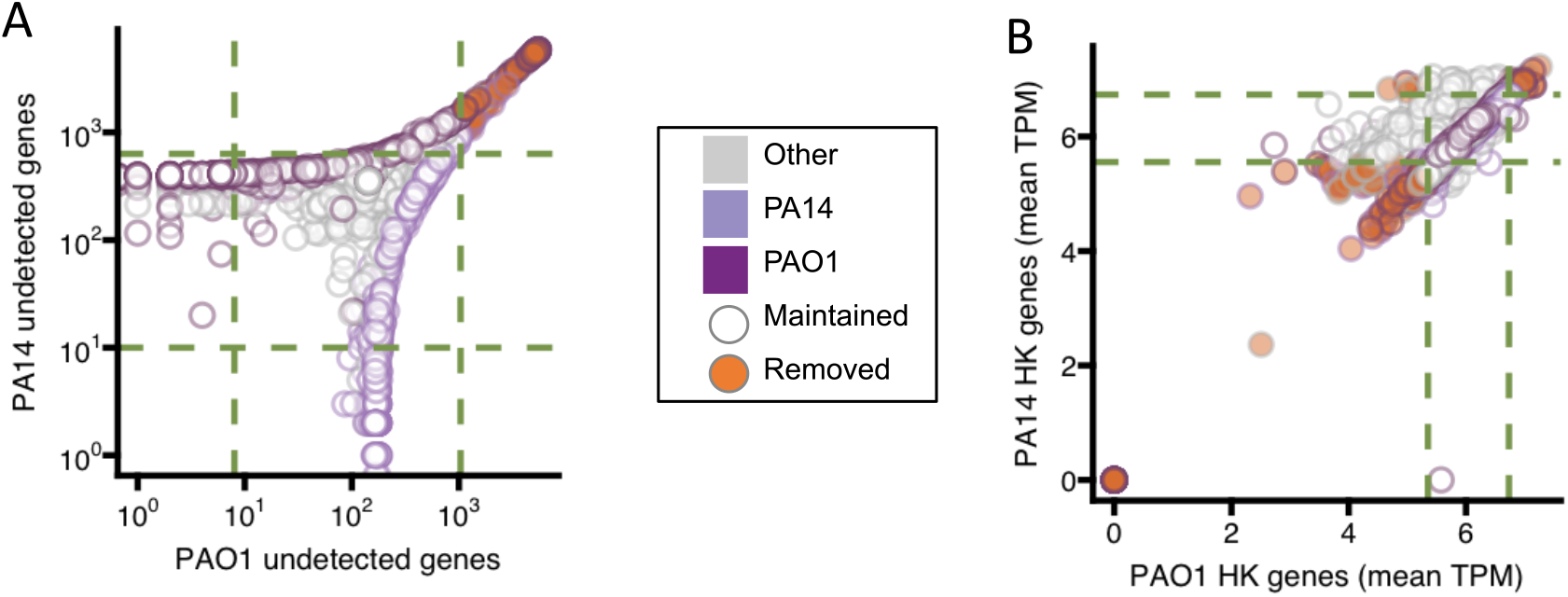
**A)** Lower and upper filtering thresholds for sparsity were defined as the 10^th^ and 90^th^ percentiles. Values for PAO1 and PA14 were separately determined using samples annotated as being each strain, indicated by circle color showing PAO1 (dark purple), PA14 (light purple) or otherwise annotated and unannotated (grey). Samples were excluded from the compendium if they fell outside the ranges by both PAO1- and PA14-defined criteria as indicated by filled circles (orange). **B)** Filtering thresholds for housekeeping gene expression were defined using the same samples as the sparsity threshold determination and as the 20^th^ and 98^th^ percentiles for PAO1 and PA14, separately determined by samples of each strain. Strain annotations and inclusion (maintained) or exclusion (removed) are indicated by circle outline and fill color as they were for sparsity filtering.

### Filtering and normalization expose correlations in known gene sets

To enable compendium-wide or multi-experiment analyses, we sought to normalize the compendium of gene expression profiles. The need for normalization was highlighted by the correlation analysis of gene expression. In a previously published compendium of microarray data (25), we observed the expected result that co-operonic genes had higher median correlation values (Pearson correlation coefficient = 0.66) than random pairs (Pearson correlation coefficient = -0.008, **Figure 5A**). Genome-wide gene-gene correlation analyses helped uncover new *P. aeruginosa* biology. However, in the RNA-seq compendium that we constructed, even random gene pairs were highly correlated (Pearson correlation coefficient = 0.42), though the average correlation for co-operonic genes was higher (Pearson correlation coefficient = 0.75). To overcome these correlations between randomly chosen gene pairs, we normalized data using the ratio of medians (RM) method, which is employed in DESeq2 (31) to account for differences in read depth (32, 33). Because it produces sample-wise normalization factors using a median-based pseudo-reference, it can be easily extended to a compendium of samples with no singular control condition. After we applied sample-wise RM normalization to the filtered PAO1-aligned and PA14-aligned compendium, correlations between randomly chosen gene pairs were essentially eliminated (−0.008) while intra-operonic correlations remained high (0.67) (**Figure 5A**).

**Figure 5.**
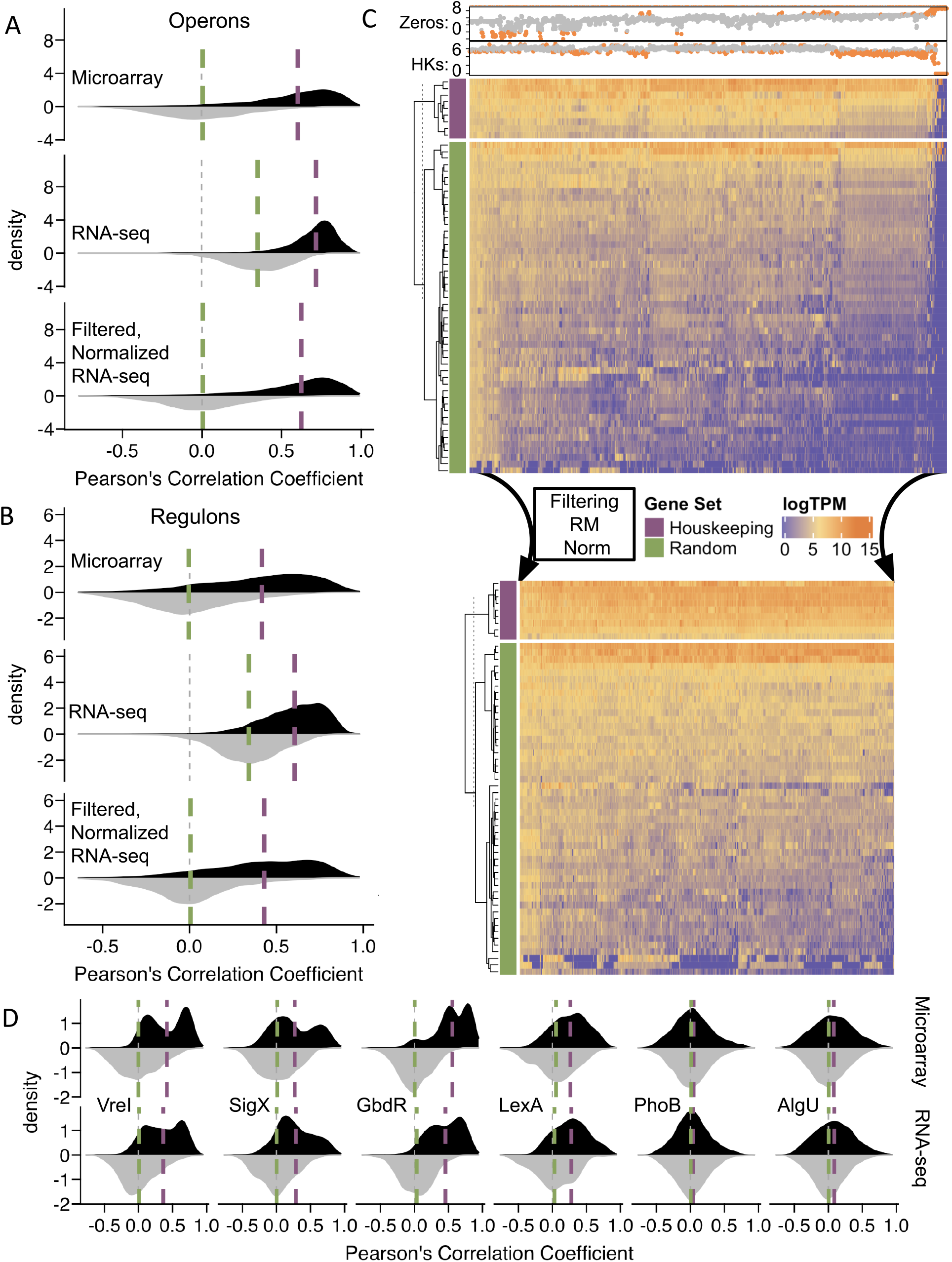
The compendium of PAO1-aligned *P. aeruginosa* gene expression profiles produced using Salmon contains patterns consistent with known transcriptional sets. **A)** Similar to a previously published array compendium, heuristic filtering and Ratio of Medians Normalization (RM) correct spurious correlations in expression of random sets of genes (grey distributions), shifting the medians (green lines) to 0 and exposing inter-operon gene-gene correlations (black distributions) with elevated medians (purple lines). **B)** Inter-regulon gene-gene correlations show a similar improvement upon filtering (removal of samples outside of zero-count (Zeros) and median housekeeping (HK) thresholds, orange points in upper annotations) and normalization compared to size-matched random controls. **C)** After filtration and median ratio normalization, samples across the compendium have more similar overall distributions of HK and 50 randomly selected (Random) genes. **D)** Within-gene set gene-gene correlation distributions are similar across the microarray and filtered, normalized RNA-seq.

Filtering and median-based normalization similarly affected correlations between genes within previously characterized, manually annotated regulons (most of which were composed of multiple operons) from the database RegPrecise (34), compared to size-matched sets of random genes (**Figure 5B**). Upon completion of the filtering and normalization steps, the marked reduction in expression correlations of randomly selected genes with each other, and with HK genes, is visually evident in a heatmap of log TPM values (**Figure 5C**). The same filtering and normalization steps applied to the compendium of reads mapped to the PA14 compendium were similarly beneficial in reducing sample-wise effects in the data (**Supplemental Figure S2**). In **Supplemental Figure S3 A and B**, we show the gene-gene correlations for operons, relative to equally sized random control groups, for each step in the creation of the PAO1-aligned and PA14-aligned compendia. We demonstrate that an alternative normalization strategy (trimmed mean of means, TMM) did not remove the sample-wise effects in the data. The complete filtered and normalized data for all profiles derived from mapping to strain PAO1 or PA14 genomes are available in **Supplemental Dataset S3** and **Supplemental Dataset S4**, respectively.

We manually curated twenty-one sets of genes regulated by transcription factors of interest from studies that deployed expression profiling analyses of mutants, DNA-binding assays, or promoter analyses (promoter fusions and motif searches). Experiments used to define gene sets were performed in a mix of strain backgrounds. The gene sets ranged in size from 5 – 405 genes, spanned multiple transcriptional units, and their gene contents were not exclusive of each other. These gene sets were involved in global biological programs such as quorum sensing, adaptation to stationary phase, metabolism of specific substrates, nutrient restriction, oxidative stress, oxygen tension, and virulence-related cues. While gene-gene expression correlations across the compendium were expectedly lower for gene sets than for operons or regulons, heuristic filtering and sample-wise normalization improved signal visibility by showing a clear elevation of median within gene set correlation (Pearson correlation coefficient = 0.26) compared to size-matched random controls (Pearson correlation coefficient = - 0.001) (**Supplemental Figure S3 C**,**D**). Some gene sets with markedly higher than random gene-gene correlations included sets of genes regulated by the extracytoplasmic sigma factors VreI and SigX and the transcription factors GbdR and LexA (**Figure 5D**). On the other hand, the gene sets regulated by the transcription factors PhoB and AlgU, did not discriminate from random control sets, and we expect that some of the hundreds of genes in these gene sets include many genes that are affected in a strain- or condition-specific manner, likely due to indirect factors. While the distribution of within-gene set gene-gene correlations varied based on gene set, some with bimodal distributions (sets of genes regulated by VreI, SigX and GdbR) and others with more normal distributions (sets of genes regulated by PhoB, AlgU and LexA), all were remarkably similar across the microarray and RNA-seq compendia (**Figure 5D**). Since the microarray and RNA-seq compendia were composed of different samples, expression relationships between genes that are reflected in both compendia suggest patterns driven by a common influence which cannot be expression profiling technology or platform, but rather is likely to be underlying biology. The effectiveness of filtering and normalization to expose compendium-wide gene-gene correlations driven by known biological mechanisms (operons, regulons and gene sets), foreshadows the potential to identify new biology based on compendium-wide correlations and provides a foundation on which future studies can rely.

### Analysis of different strains, media, treatments and genetic perturbations profiled in the compendium

Curation of metadata is a valuable step in compendium creation because it enables users to investigate trends associated with treatment conditions or strains across multiple studies. When constructing our previously published microarray compendium of *P. aeruginosa* gene expression data, metadata was manually collected and curated by experts (26). However, because of the time-consuming nature of manual metadata curation, it is a process difficult to scale to larger compendia such as the RNA-seq compendium presented here. To meet the challenge of providing curated metadata for the RNA-seq compendium, we employed an R package called GEOquery that automates collection of metadata associated with studies present in the Gene Expression Omnibus (GEO). Of the 277 BioProjects contained in the compendium, about half (139) are present in GEO and therefore have documented metadata amenable to automated parsing (see **Supplemental Dataset S5**). Using these metadata, we analyzed the composition of the compendium with respect to *P. aeruginosa* strains. The annotated compendium contains approximately 70 studies that used strain PAO1 (∼700 total profiles) and 30 studies that used strain PA14 (∼400 total profiles), while the remaining studies used other laboratory strains (PAK, PA3) or clinical and environmental *P. aeruginosa* isolates (630 profiles) (**Figure 6A**). As an alternative approach to identifying the strain for each sample via metadata, we also used the filtered and normalized compendia themselves to classify samples as PAO1-like or PA14-like using accessory genes that were present in only one of these strains and did not have homologs in the other strain. For each sample, the median expression of PAO1- and PA14 strain-specific accessory genes was calculated. The profiles that were annotated as either being PAO1-like or PA14-like in the SRA database were validated using this approach (**Figure 6B**). Strain PAK, which is more similar to PAO1, was also correctly assigned using this approach. Samples with high counts for a mixture of PAO1- and PA14-specific accessory genes belonged to clinical isolates or samples without strain annotations (referred to as Other in **Figure 6B**). We further assessed strain assignment using the bimodal distributions of median expression of PAO1- and PA14-specific accessory genes which accurately separated strains with known annotations (**Supplemental Figure S4 A**,**B**). In **Supplemental Dataset S2**, the median accessory gene set value is reported for both PAO1 and PA14 accessory genes in all samples in the filtered, normalized PAO1-aligned and PA14-aligned compendia.

**Figure 6.**
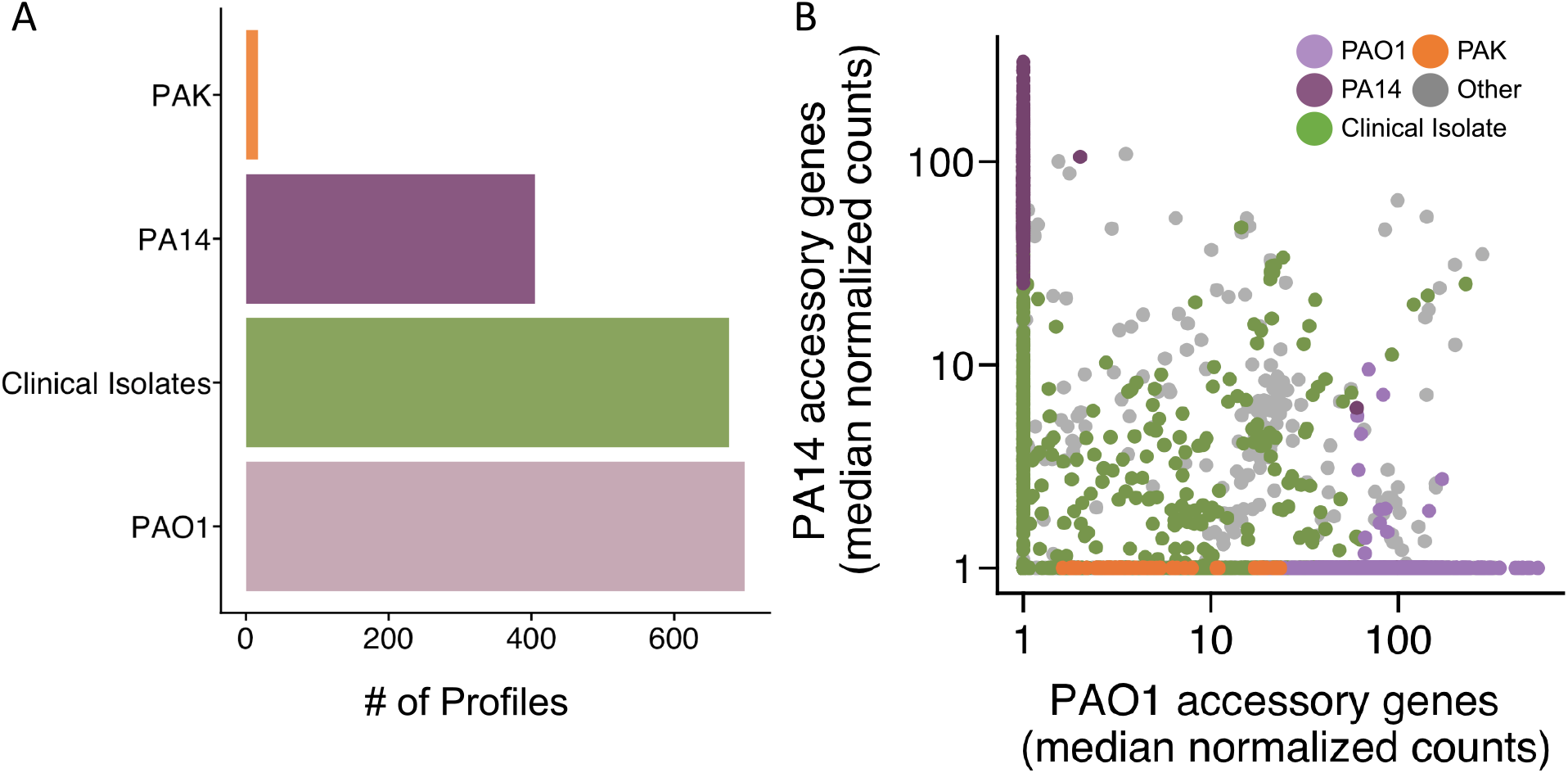
Researchers have interrogated a variety of different *P. aeruginosa* strains and PAO1- or PA14-likeness can be determined by expression levels of PAO1- and PA14-only genes. **A)** As determined by annotations, PAO1 is the most commonly-studied strain, employed in over 70 studies in a total of nearly 700 profiles. **B)** In the absence of annotations, samples can be binned into PAO1 or PA14 categories based on the expression values of PAO1-specific and PA14-specific accessory genes. Samples annotated as anything other than PAO1, PA14, PAK or clinical isolates (including samples without any strain annotation) were grouped as “Other”.

Analysis of clinical isolates for median expression of PAO1-specific or PA14-specific accessory genes (**Supplemental Figure S4C and D**) found that the median expression for either set was generally lower than for the authentic PAO1 or PA14 samples. Examples of profiles that have relatively high PAO1 accessory and PA14 accessory expression included two profiles from clinical isolates (SRX1096902, SRX1097013). Such samples suggest new opportunities for analyzing the expression of specific accessory genes across diverse strain background using normalized compendia such as those reported here. If necessary, further optimization of mapping sensitivity and using a chimeric or decoy transcriptome indices could remove erroneous mapping. This data-driven approach for meta-data assignment can easily extend to new datasets and new parameters that reflect specific conditions.

### Other compendium characteristics can be analyzed from GEO metadata

To gain more insight into biases in experimental design that might influence gene expression patterns identified in the compendium, we used metadata gathered from the Gene Expression Omnibus (GEO) that was cleaned and curated (**Figure 7A**). These metadata revealed that studies in the compendium also employed a wide variety of different media. Approximately 70 studies - and nearly 1,250 total profiles - were conducted with LB medium (which includes annotations of lysogeny broth, Lennox broth, Luria-Bertani medium) which was the most common medium category (**Figure 7B**). Approximately half of PAO1 studies and half of PA14 studies used LB medium; clinical isolates were studied predominantly in LB. Other media types include minimal media, such as M9 minimal medium, and complex media, such as synthetic cystic fibrosis medium (SCFM and SCFM2) (35, 36) (**Figure 7B**). Some media were only studied in one strain. For example, M9 medium was only used in PAO1 studies, while Mueller-Hinton Broth was exclusive to PA14 studies (**Figure 7C**). These analyses highlight that analysis of media effects should include consideration of strain when there is a bias. A more detailed curation of media conditions across individual experiments will expand the potential to develop hypotheses based on expression trends.

**Figure 7.**
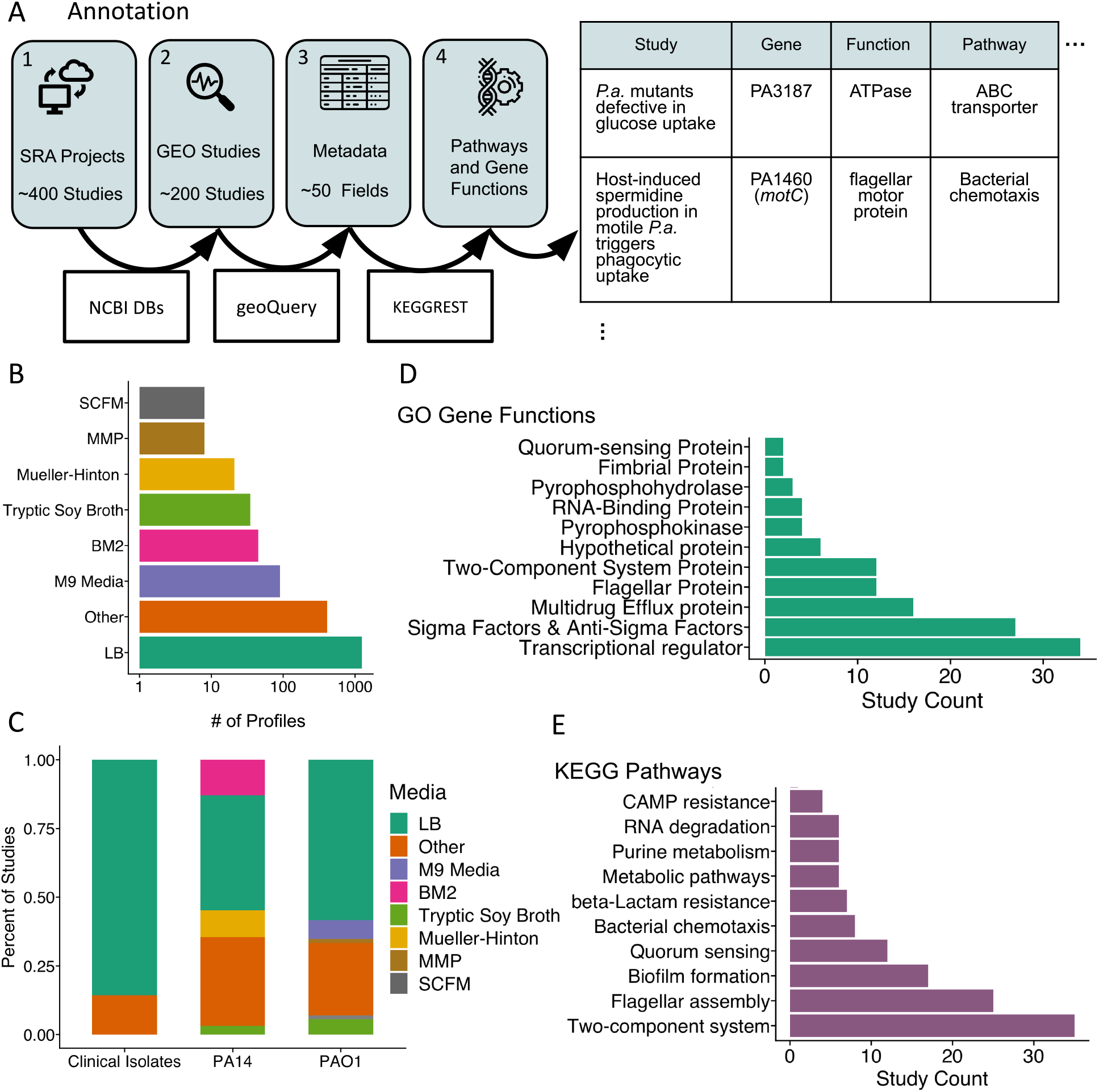
Researchers have interrogated a variety of different *P. aeruginosa* strains in a range of different media and a variety of genes have been knocked-out or over-expressed. **A)** Metadata was gathered from the Gene Expression Omnibus (GEO), and was cleaned and curated prior to extension of gene annotations to pathways and functions. **B)** LB (lysogeny broth, Lennox broth, and Luria Bertani medium) is the most common medium used for samples in the compendium. **C)** The two most common *P. aeruginosa* laboratory strains have been studied in various media, and clinical isolates have been predominantly studied in LB. **D)** Of the biological functions frequently under investigation, transcriptional regulators were most commonly manipulated, followed by sigma factor or anti-sigma factor genes and multidrug efflux pump genes. **E)** Manipulated genes belong to various KEGG pathways including multiple genes and studies investigating two-component systems. In 7D and 7E, a single gene contributes to the study count of multiple pathways if the gene is found on more than one functional classification or pathway.

In a number of studies, researchers had performed some form of genetic perturbation - the knockout or overexpression of a gene or genes - then interrogated the effects of this manipulation on *P. aeruginosa* gene expression. Among the GEO studies in which researchers performed genetic perturbations and identified the genes that they perturbed, there was a notable emphasis on regulators (e.g. transcription factors, sigma factors and two component systems) among the associated GO terms (**Figure 7D**) and KEGG pathways (**Figure 7E**). Other functional categories include appendages, biofilm, quorum sensing, drug resistance and metabolic pathways; an important caveat to keep in mind is that annotations may be incomplete or may reflect indirect or condition-specific effects (**Supplemental Dataset S6**).

## Conclusion

There are multiple microbes for which there are thousands of publicly available transcriptome datasets including *Escherichia coli, Staphylococcus aureus, Saccharomyces cerevisiae* and *Candida albicans*. Here, we present the largest collection of publicly available *P. aeruginosa* gene expression data with twice as many samples as are in our previously published microarray compendium. Further, this collection is an order of magnitude larger than any other collection of RNA-seq data made available to the community (37). The analysis of these data, which include diverse experiments, strains, medium conditions and mutants, can greatly aid the generation of hypotheses that can be rigorously tested by more targeted analysis of new or existing datasets and orthogonal experimental approaches. Our previous studies using a compendium of *P. aeruginosa* microarray data highlight the value of compendium-wide analyses to describe robust regulons which can aid in the development of biomarkers for analysis of pathway activities in clinical or environmental samples. Furthermore, the availability of a filtered, normalized and annotated compendium will enable systems-levels analyses of pathway-pathway relationships. The workflow presented here is scalable and adaptable to new datasets and organisms and provides a critical approach to fully utilize transcriptomics data which is maintained in databases for re-analysis but is too often left waiting.

## Methods

### RNAseq processing

The RNA-seq dataset used to compare CLC and Salmon alignments consisted of wild type PA14 (DH122) and *pstB*::Tn*M* (DH753) grown as colony biofilms on 3-(N-morpholino) propanesulfonic acid (MOPS) 0.2% glucose, 0.7 mM phosphate plates with 1.5% agar, in duplicate. Cells were collected as cores from agar plates: cores were taken using a straw, cells were suspended by shaking agar plugs in 1 mL dH_2_O on the Disrupter Genie for three minutes. RNA was isolated using the Qiagen RNeasy kit (cat. 74004) and DNase treated using the Turbo kit (cat. AM2239), libraries were prepped in accordance with Illumina protocols, including ribo-depletion, and sequenced on the Illumina NextSeq at the Geisel School of Medicine at Dartmouth in the Genomics Shared Resource. RNA-seq data was processed using the CLC Genomics Workbench v10 and differential expression analysis was performed using EdgeR (23).

### Salmon Mapping

Salmon was run in mapping-based mode to make use of its fast mapping algorithm. To create the transcriptome indices from the reference genomes, the Salmon index call was used with 15 set as the minimum length for an acceptable alignment match (k=15). To align reads, the Salmon quant call was used with the validation method set to ‘score’ and ‘validate mappings’. Code is available at github.com/hoganlab-dartmouth/pa-seq-compendia along with all generated data.

### Determination of heuristic criteria

Housekeeping genes were chosen from the literature and *P. aeruginosa* gene expression field standards (39, 40). Percentile cutoffs were based on visual inspection and the removal of datasets with technical differences (meta-transcriptomic data and RIP-seq data).

### Creating separate PAO1 and PA14 compendia

After the filtering heuristic step, there were 2,333 samples mapped to 5,563 genes using the PAO1 reference genome and 5,891 genes using the PA14 reference genome. Although SRA provides annotations for strain type, we used the expression activity of the accessory genes (i.e. genes that are specific to PAO1 or specific to PA14 according to the *Pseudomonas* Genome Database ortholog annotations) to bin samples into PAO1 and PA14 compendia.

### Compendium Normalization

Transcripts per million (TPM) were estimated by Salmon and exported directly. Trimmed mean of means (TMM) and ratio of medians (RM) normalization was applied to the estimated counts exported from Salmon using the R packages EdgeR (23) and DEseq2 (31), respectively. For both normalization methods, per-sample coefficients were extracted and multiplied by estimated counts. After normalization the compendia were 0-1 scaled with a linear transformation based on the matrix maximums and minimums. The array compendium was downloaded from the ADAGE github repository (https://github.com/greenelab/adage) and had already been 0-1 scaled with linear transformations based on gene-wise maximums and minimums.

### Correlation Analyses

Pearson correlation coefficients were calculated in R using the base corr function on predetermined sets of genes (operons from the Pseudomonas Genome Database, regulons from RegPrecise, gene sets from select publications) and averages of size-matched sets of randomly selected genes (n=10). Distributions and medians were plotted in ggplot2 (41).

### Annotations

Detailed metadata were gathered for all the *P. aeruginosa* RNA-seq experiments present in the Gene Expression Omnibus (GEO) with select fields: strain, media, genetic perturbation, and other experimental conditions. *P. aeruginosa* RNA-seq studies were identified in GEO by searching for ‘Pseudomonas aeruginosa’ in the GEO Data Set (GDS) browser, filtering by the study type ‘Expression profiling by high throughput sequencing’ (which corresponds to RNA-seq studies), and further filtering by organism to select only studies containing *P. aeruginosa* RNA-seq data (i.e., not RNA-seq studies of mice or human cells exposed to *P. aeruginosa*, which came up with the basic search).

Metadata for each *P. aeruginosa* RNA-seq study was downloaded directly from GEO as a .txt file. This summary contained an FTP download link for each study, in which the GSE accession for the study was embedded. For all studies, the accession was extracted with the str_extract_all() function from the R package stringr, and was subsequently fed into the function getGEO() of the GEOquery R package from Bioconductor (42). The getGEO() output is a large R list object containing very detailed metadata on the strains, media, treatment, and other conditions employed for each study.

Metadata was further cleaned in Excel so that the metadata could be re-uploaded to R in a suitable format for figure creation. All figures involving annotation data were created with the R package ggplot2. Additional information was gathered from the Kyoto Encyclopedia of Genes and Genomes (KEGG) Rest server using the package KEGGREST from Bioconductor (43). For *P. aeruginosa* RNA-seq studies that perturbed certain genes, the common gene names provided in the metadata (e.g., *sigX*) were converted manually to their respective locus tags (e.g., PA1776) and fed into KEGGREST’s keggGet() function, which extracted information on associated KEGG pathways and gene functions.

## Data availability

All processed data and necessary code to recapitulate analyses are available on the pa-seq-compendia github repository (github.com/hoganlab-dartmouth/pa-seq-compendia) and raw data for the original RNA-seq analyses presented in this study are available on the NCBI SRA database and Gene Expression Omnibus under the accession GSE192694.

## Acknowledgments

Research reported in this publication was supported by grants from the Cystic Fibrosis Foundation HOGAN19G0 (D.A.H.), GREENE21GO (C.S.G) and STANTO19R0 (B.A.S) and the National Institutes of Health (NIH) through NIDDK P30-DK117469 (B.A.S. and S.N.) and R01 HL151385 (B.A.S.). Additional core facility support came from the NIH NIGMS P20GM113132 (BioMT). The content is solely the responsibility of the authors and does not necessarily represent the official views of the NIH. The funders had no role in study design, data collection and analysis, decision to publish, or preparation of the manuscript.

## Supplemental Figure Legends

**Supplemental Figure S1. Salmon parameter choice of unmapped reads**. A comparison of transcripts per million (TPM) for a single *P. aeruginosa* strain PA14 RNA-seq sample (WT, rep 1) mapped by the aligner CLC to TPMs determined by Salmon in mapping mode with the library type specified as either paired end (paired, orange) or single end (unpaired, purple). When the library type was specified as “paired”, Salmon underestimated some transcripts compared to CLC.

**Supplemental Figure S2. Heuristic filtering and Ratio of Medians Normalization (RM) correct spurious correlations in expression of a random set of genes and housekeeping genes in a PA14-mapped compendium. A)** PA14-aligned compendia after filtration (removal of samples outside of zero-count (orange points in Zeros panel) and median housekeeping (HK) thresholds, orange points in HKs panel), and RM normalization have more similar overall distributions of housekeeping and 50 randomly selected genes (Random).

**Supplemental Figure S3. Effects of different steps in the creation of the compendium on the gene-gene correlations within operons and gene sets. A)** The PAO1-aligned compendium, similar to a microarray compendium, can be filtered and normalized to correct spurious correlations in expression of random sets of genes (grey distributions), shifting the medians (green lines) to 0 and exposing intra-operon gene-gene correlations (black distributions) with elevated medians (purple lines). TPM, transcripts per million; Filtered TPM, after filtering steps outlined in Fig. 5C; TMM, Trimmed mean of means; RM, ratio of medians normalization. **B)** The PA14-aligned compendium responds to filtering and normalization similarly to the PAO1-aligned compendium with respect to operons. **C)** The PAO1-aligned compendium responds to filtering with respect to gene sets similarly as with respect to operons. **D)** The PA14-aligned compendium responds to filtering and normalization similarly to the PAO1-aligned compendium with respect to gene sets.

**Supplemental Figure S4. In the absence of annotations, profiles can be binned into PAO1 or PA14 categories based on the expression values of PAO1-specific and PA14-specific accessory genes. A)** Samples known to be of strain PAO1 had profiles with higher median expression of PAO1 accessory genes (PAO1) than did non-PAO1 profiles including PA14, PAK, clinical isolates and profiles without strain annotations (Other). **B)** Samples known to be of strain PA14 had higher median expression of PA14 accessory genes (PA14) than did non-PA14 profiles (Other). **C)** Clinical isolates had lower median expression of PAO1 accessory genes relative to strains that are not annotated as clinical isolates (Other). **D)** Clinical isolates had low median expression of PA14 accessory genes relative to strains that are not annotated as clinical isolates (Other) samples.

## Supplemental Dataset Legends

**Supplemental Dataset S1: WT *pstB* differential expression analyses**.

Comparison of CLC full alignment and salmon mapping (pseudo-alignment) pre-processing effects on downstream differential expression analysis comparing WT PA14 *P. aeruginosa* gown in sufficient phosphate to suppress the low-phosphate response and the mutant derivative with a disrupted repressor, *pstB*::Tn*M*, and thus a constitutive low-phosphate response in these conditions. **(Columns A-D)** Annotations of PA14 locus tags, PAO1 homologs, gene names and whether each gene is regulated by PhoB in response to low phosphate. (**Columns E-S)** EdgeR differential expression analysis results for WT vs. *pstB*::Tn*M* were aligned to the PA14 reference genome in CLC and salmon and aligned to the PAO1 reference genome in Salmon. RNA extracted from colony biofilms on MOPS 0.7 mM Pi, 0.2% glucose. **(Columns T-AE)** Raw counts values for WT and *pstB*::Tn*M* samples, used in the differential expression analyses. (Columns AF-AQ) TPM values for WT and *pstB*::Tn*M* samples used in the differential expression analyses.

**Supplemental Dataset S2: Experiment Profile Characteristics**

Filter criteria (housekeeping gene expression and zeros counts) values, accessory gene expression median values and experiment (profile) - wise annotations for the pre-filtered compendium.

**Supplemental Dataset S3: PAO1-aligned compendium**. Filtered and normalized PAO1-aligned compendium. Please note this file is truncated to two decimal points to reduce the file size. An untruncated version is available hosted by the Center for Open Science under the *P. aeruginosa* RNA-seq compendia project (https://osf.io/s9gyu/).

**Supplemental Dataset S4: PA14-aligned compendium**. Filtered and normalized PA14-aligned compendium. Please note this file is truncated to two decimal points to reduce the file size. An untruncated version is available hosted by the Center for Open Science under the *P. aeruginosa* RNA-seq compendia project (https://osf.io/s9gyu/).

**Supplemental Dataset S5: Full Annotations By Study**.

Annotations from studies in GEO that make up ∼50% of the total compendium. Annotations collected for studies in GEO including all compiled fields.

**Supplemental Dataset S6: Full Functional Annotations**.

KEGG pathway and GO molecular function annotations of studies with perturbed gene expression.

